# Structural mechanism for recognition of E2F1 by the ubiquitin ligase adaptor Cyclin F

**DOI:** 10.1101/2025.01.15.633208

**Authors:** Peter Ngoi, Xianxi Wang, Sivasankar Putta, Ricardo F. Da Luz, Vitor Hugo B. Serrão, Michael J. Emanuele, Seth M. Rubin

## Abstract

Cyclin F, a non-canonical member of the cyclin protein family, plays a critical role in regulating the precise transitions of cell-cycle events. Unlike canonical cyclins, which bind and activate cyclin-dependent kinases (CDKs), Cyclin F functions as a substrate receptor protein within the Skp1-Cullin-F box (SCF) E3 ubiquitin ligase complex, enabling the ubiquitylation of target proteins. The structural features that distinguish Cyclin F as a ligase adaptor and the mechanisms underlying its selective substrate recruitment over Cyclin A, which functions in complex with CDK2 at a similar time in the cell cycle, remain largely unexplored. We utilized single-particle cryo-electron microscopy to elucidate the structure of a Cyclin F-Skp1 complex bound to an E2F1 peptide. The structure and biochemical analysis reveal important differences in the substrate-binding site of Cyclin F compared to Cyclin A. Our findings expand on the canonical cyclin-binding motif (Cy or RxL) and highlight the importance of electrostatics at the E2F1 binding interface, which varies for Cyclin F and Cyclin A. Our results advance our understanding of E2F1 regulation and may inform the development of inhibitors targeting Cyclin F.

## Introduction

Cyclin F is a non-canonical member of the cyclin family and is distinguished by its lack of a direct role in CDK activation. Instead, Cyclin F functions as a substrate receptor for the SCF (Skp1-Cullin-F-box) E3 ubiquitin ligase complex (1, 2) (Figure 1A), facilitating the ubiquitylation and subsequent degradation of key proteins involved in cell cycle regulation (3, 4), DNA replication (5, 6), and the DNA damage response (7, 8). The expression profile of Cyclin F is similar to the canonical cell-cycle and CDK-activating Cyclin A, with mRNA Cyclin F accumulation beginning in early S phase, peaking in G2, and declining before mitosis (9). Notable targets of Cyclin F include E2F transcription factors, the E2F repressor p130, RRM2, and CPP110, which are crucial for controlling cell proliferation, maintaining genomic stability, and responding to DNA damage (3–8). Disruption of Cyclin F function and regulation has been linked to various phenotypes, including cell-cycle disruption and genomic instability, and has implications in cancer development and neurodegenerative diseases (2, 10). Interestingly, emerging evidence also suggests Cyclin F may play a role in neurodegenerative diseases due to its involvement in degrading proteins such as TDP-43 and HSP90, which are implicated in amyotrophic lateral sclerosis (ALS) (11, 12). The role of Cyclin F in disease and in maintaining cellular homeostasis highlights its broad impact in human health and serves as motivation to investigate the structural mechanism of substrate recruitment.

**Figure 1.**
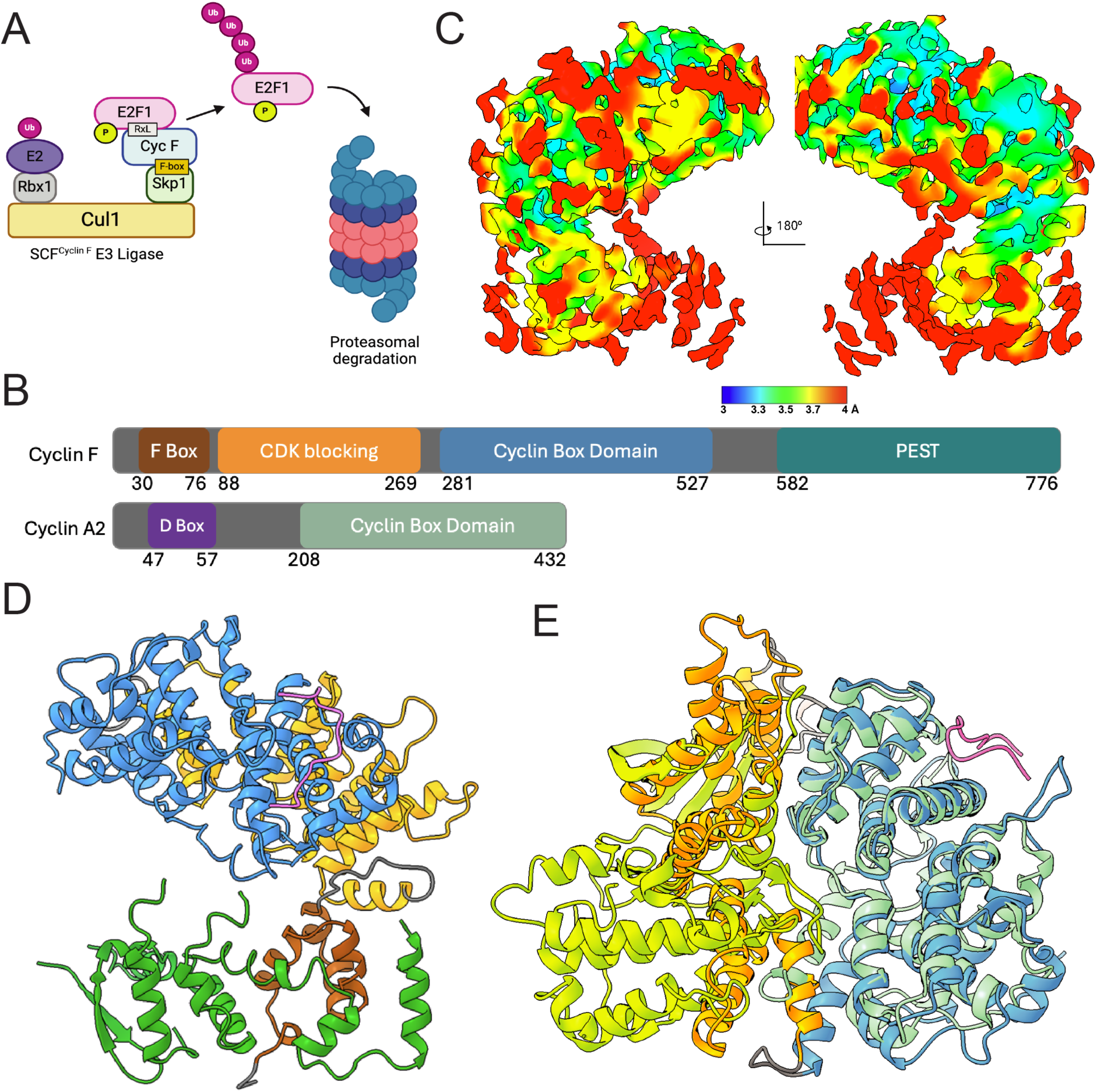
**A)** Model for E2F1 ubiquitylation by SCF^CyclinF^ and proteasomal degradation. Cyclin F binds and activates the Skp1-Cul1-F box (SCF) E3 ligase to E2F1 and other substrates. **B)** Domain architecture for Cyclin F and Cyclin A. **C)** Final electron density map of E2F1-Cyclin F-Skp1 complex with local resolution from 3.0-4.0 Å, visualized by the color-coded surface. **D)** Final model of the E2F1-Cyclin F-Skp1 complex (PDB: 9CB3). E2F1 (pink), Skp1 (green), and Cyclin F (domains colored according to figure B) are shown. **E)** Overlay of E2F1-Cyclin F (9CB3) and E2F1-CDK2-Cyclin A (PDB: 1H24) cyclin domains. E2F1 peptide (pink), Cyclin A (green), CDK2 (yellow), Cyclin F cyclin domain (blue), and Cyclin F CBD (orange) are shown.

The regulation of the cell cycle is a fundamental process that ensures the precise replication and distribution of genetic material in eukaryotic cells. Central to this regulation are the E2F family transcription factors, particularly E2F1-3a (called hereafter E2F), which drive the expression of genes essential for DNA synthesis during S phase (13–16). E2F activity, which requires the formation of a heterodimer with a DP family protein, must be tightly controlled. Until S phase, E2F is bound and inhibited by the retinoblastoma protein (RB); RB phosphorylation releases E2F leading to its activation (17). Once DNA replication is complete, E2F must be downregulated to facilitate the transition out of S phase and prevent unscheduled re-initiation of DNA replication. This downregulation is achieved through phosphorylation by the CDK2-Cyclin A complex, which inhibits E2F-DNA binding (18–20), and subsequent degradation via the ubiquitin-proteasome system (UPS) (21). Cyclin F has been identified as a key protein for the degradation of E2F (3, 21, 22). The interaction between Cyclin F and E2F is of particular interest, as misregulation of E2F activity is a hallmark of tumorigenesis, and manipulation of the interaction between Cyclin A and E2F is being explored as a therapeutic strategy (23–25).

The substrate adaptor protein in SCF family ubiquitin ligases associates with Skp1 and binds substrate to recruit it to the complex (Figure 1A). Cyclin F contains the predicted key substrate adaptor structural domains (Figure 1B, Supplemental Figure 1), including the F-box domain (residues 30-76) that facilitates Skp1 binding. The cyclin box domain (residues 281-587) is conserved among all canonical cell-cycle cyclin proteins. Cyclin F also contains a C-terminal PEST domain (residues 582-776), which is predicted to be disordered and is involved in regulating stability (9). Additionally, Cyclin F possesses a predicted helical domain that is unique among cyclins (residues 99-269). This Cyclin F-specific domain is located between the F box and cyclin domains and has an undefined function.

Several previous studies have shown the importance of an RxL motif within CDK substrates, including E2F, RB family proteins, and CIP/KIP inhibitors such as p21 (26–29). This short linear interacting motif (SLIM) binds the cyclin box domain at a site that contains the conserved hydrophobic groove, which is called the MRAIL site after the sequence that helps form the groove in Cyclin A. Observations suggest that there are sequence elements beyond the conserved minimal RxL and MRAIL motifs that influence the selective recruitment of cyclin substrates and inhibitors. For example, the inclusion of residues N-terminal to the RxL increases the affinity of peptides for Cyclin A (28), p27 binds to CDK2-Cyclin A and CDK2-Cyclin E with different affinities (30), and replacing an RxL-containing sequence in E2F4 with the RxL sequence from E2F1 promotes E2F4 phosphorylation by CDK2-Cyclin A (31). Cyclin F has also been reported to necessarily recruit substrates through their RxL motifs, which are likely to form direct contacts with a Cyclin F-MRAIL motif (MRYIL in Cyclin F) for ubiquitin-mediated degradation (3, 4). Several important questions surrounding substrate recognition by cyclins remain, including to what extent the sets of Cyclin F and Cyclin A substrates overlap, what are the determinants of specificity in substrate recognition, and what structural differences among cyclin proteins account for variations in substrate recognition.

Here, we present the structure of an E2F1-Cyclin F-Skp1 complex that we determined using single-particle cryo-electron microscopy (cryo-EM). The structure and additional fluorescence polarization (FP) affinity measurements highlight the unique features of Cyclin F that distinguish it from Cyclin A, particularly the inability of Cyclin F to bind and activate CDKs and to interact differently with RxL substrates. We identified Cyclin F-specific residues, outside of its canonical MRYIL motif, that impact substrate binding, enhancing our understanding of the regulatory mechanisms governing Cyclin F activity. While the hydrophobic MRYIL motif has been shown to be necessary for binding, our evidence highlights the importance of electrostatic interactions with RxL SLIMs that affect Cyclin F and Cyclin A differently. These data suggest mechanisms by which Cyclin F might bind to some, but not all, Cyclin A targets, guiding our understanding of whether substrates are either phosphorylated or ubiquitinated. This research could inform the development of new inhibitors targeting Cyclin F and E2F.

## Results

### Single particle cryo-EM reveals the structure of Cyclin F

We employed single-particle cryo-EM to determine the structure of Cyclin F and the molecular details of the E2F binding interface (Supplemental Table 1, Supplemental Figure 2). Human Cyclin F (residues 1-636) and Skp1 (full length, residues 1-160), were co-expressed in sf9 insect cells, and a synthetic E2F1 peptide containing the RxL motif (GRPPVK<u>RRL</u>DLETDHQ) was added at a 2:1 molar ratio prior to vitrification. The cryo-EM data resulted in a final reconstructed map at 3.5 Å overall resolution and with local resolution spanning from 3.0 to 4.0 Å (Figure 1C and Supplemental Figures 3 and 4). A final E2F1-Cyclin F-Skp1 model was built from a starting predicted model of the Cyclin F structure (Figure 1D, Supplemental Figure 4).

The Cyclin F structural model is composed of three alpha-helical domains: the F box, the cyclin box, and the Cyclin F-specific helical domain that we named the CDK blocking domain (CBD) (Figure 1D). Amino acid sequences C-terminal to the cyclin box domain were included in the protein construct used for EM studies but could not be modeled; the lack of corresponding electron density at the adopted contour level of the map is consistent with the PEST domain being intrinsically disordered. The F box domain binds Skp1 similarly to that observed in other F box-Skp1 protein complexes, including the Skp2-Skp1 and FBXW7-Skp1 complex (Supplemental Figure 5A and 5B), a necessary interaction for assembling the substrate adaptor subunit with the rest of the SCF E3 ubiquitin ligase complex (Supplemental Figure 5C) (32–34).

The F box domain is followed in sequence by the CBD. The two domains are structurally connected through hydrophobic interactions involving alpha helix 5 of the CBD (α5), a 12 amino-acid loop bridging the CBD and F box domains (between α4 and α5), and alpha helices of the F box domain (α2 and α4) (Supplemental Figure 5D). Specifically, L90 on α5 interacts with F78 on L1 and V49 and L53 on α2, while L81 on L1 engages with V49 on α2. Together these residues form a hydrophobic core that fixes the orientation of the CBD and cyclin box relative to the F box and, considering the known rigid interactions between Skp1 and Cul1, ultimately the rest of the E3 ligase.

The CBD is composed of 10 alpha helices (α5-α14) that interface with alpha helices α15, α19, α21, and α22 of the cyclin box domain (Supplemental Figure 6). The cyclin box in Cyclin A binds CDK2 through an extensive (∼3550 Å^2^) interface that uses a surface opposite the MRAIL motif. Notably, the equivalent CDK-binding region in Cyclin F is inaccessible due to the interactions with the CBD, which motivates us to refer to it as the CDK-blocking domain. There are two loops in the CBD (between α6 and α7 and between α8 and α9) that form a pocket with α19 and α21 in the cyclin domain; this pocket has been implicated as a secondary substrate binding site for specific substrates (20).

### Structural details of the Cyclin F-E2F1 binding interface

The structure of the Cyclin F cyclin box aligns well with the structure of other cyclins, including Cyclin A (Supplemental Figure 7A). Cyclin F interacts with the E2F1 peptide via alpha helices α16, α18, and α20 (Figure 2A and Supplemental Figure 7B). The peptide spans ∼20 Å and makes a ∼90° turn at the C-terminal end of α16 away from α20. Several aspects of this interface are highly similar to the E2F1-Cyclin A complex. The hydrophobic groove (composed of MRYIL in Cyclin F or MRAIL in Cyclin A) is located at the N-terminal end of α16 and provides two hydrophobic pockets that bind the leucine in the RxL motif (L92 in E2F1) and an additional leucine in the +2 position from the motif (L94) (Supplemental Figure 8A and 8B). The arginine in the motif (R90) makes a salt bridge with E319 in α16 of Cyclin F, which also resembles a similar interaction made with E220 of Cyclin A.

**Figure 2.**
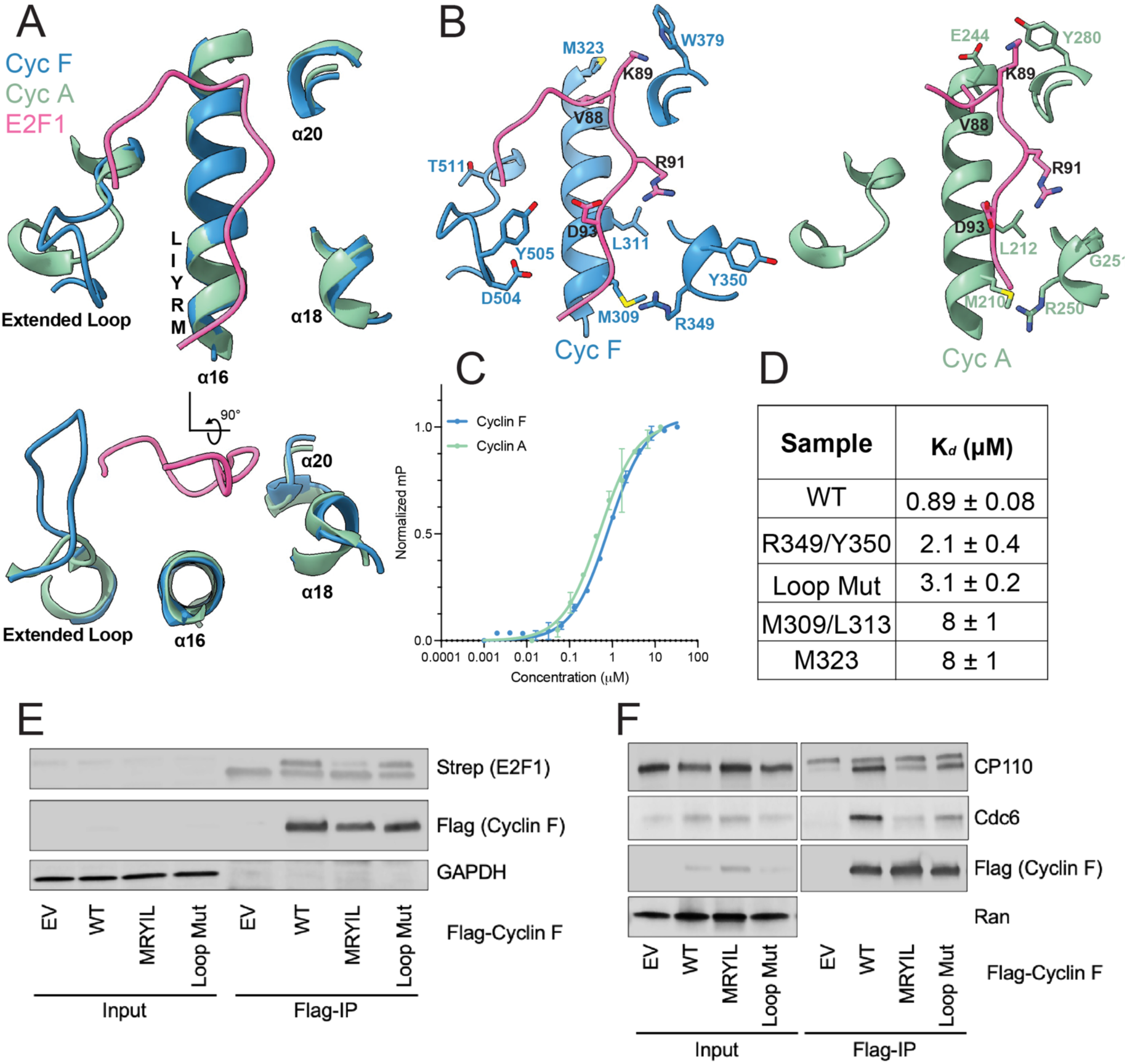
**A)** Overlay of the E2F1-binding interface in Cyclin F and Cyclin A. The E2F1 peptide is from the E2F1-Cyclin F structure. **B)** Location of amino acids that are not conserved between Cyclin F and Cyclin A are rendered as sticks. **C)** Fluorescence polarization (FP) binding curves for Cyclin F-Skp1-GST and GST-Cyclin A for the TAMRA-E2F1 peptide. **D)** FP binding affinity values of alanine mutant Cyclin F proteins for TAMRA-E2F1 peptide. The Loop Mut contains alanine substitutions in the extended loop at all residues between D504-T511. **E)** HEK293T cells were co-transfected with plasmids encoding Flag-Cyclin F wild type (WT), a mutant M309A/L311A (MRYIL) mutant, or a mutant containing D504-T511 all changed to alanine (Loop Mut) along with a plasmid encoding Strep-E2F1. Anti-Flag antibodies were used for coimmunoprecipitation, and Western blots were performed with anti-Strep and anti-Flag antibodies to detect Strep-E2F1 and Flag-Cyclin F, respectively. **F)** HEK293T cells were transfected with plasmids encoding Flag-Cyclin F WT, the MRYIL, or Loop Mut. Coimmunoprecipitation with anti-Flag antibodies was followed by Western blot analysis to detect endogenous CP110 and Cdc6.

We turned our attention to non-conserved residues around the MRYIL site that may contribute to Cyclin F substrate specificity. By comparing the E2F1-binding site in Cyclin F with Cyclin A (28), we identified key differences. Cyclin F contains a loop between α27 and α28 (residues D504-T511) that lies close to α16 and is within proximity to interact with the E2F1 peptide (Figure 2A). Notably, this loop is a region of structural divergence between the canonical cyclin proteins and Cyclin F, and the loop is longer in Cyclin F than in Cyclin A, so we call it the extended loop (Supplemental Figures 1 and 7B). The extended loop is likely flexible considering the weak electron density that is visible for the loop side chains; however, the main chain density supports its position adjacent to α16 (Supplemental Figure 3). Multiple Cyclin F side chains can be modeled within 3-5 Å of the E2F1 peptide, including D504, Y505, and T511 (Figure 2B). T511 is reported to be phosphorylated by VRK1 kinase (www.phosphosite.org) and is a VRK1 kinase consensus site; however, our data does not support phosphorylation of the recombinant protein at this site.

Around the E2F1 binding site toward the C-terminal end of α16, there are a number of non-conserved residues between Cyclin F and Cyclin A. M323 in Cyclin F is 3.5 Å away from V88 in the E2F1 peptide and appears to contribute van der Waals interactions (Figure 2B). In Cyclin A, E224 is located in this position and, alternatively, contributes to an electronegative patch in Cyclin A (Figure 3). Another site of divergence is located adjacent to the MRYIL motif in α18 in the cyclin box (Figure 2B), where Y350 is within proximity to interact with R91 of the E2F1 peptide. While electron density suggests Y350 is directed away from the E2F1 peptide (Supplemental Figure 9), the sidechain proximity and flexibility may allow for some degree of hydrogen bonding with R91. In the E2F1-Cyclin A structure, G251 is located at this position and likely does not make any contributions to binding the E2F1 SLIM.

**Figure 3.**
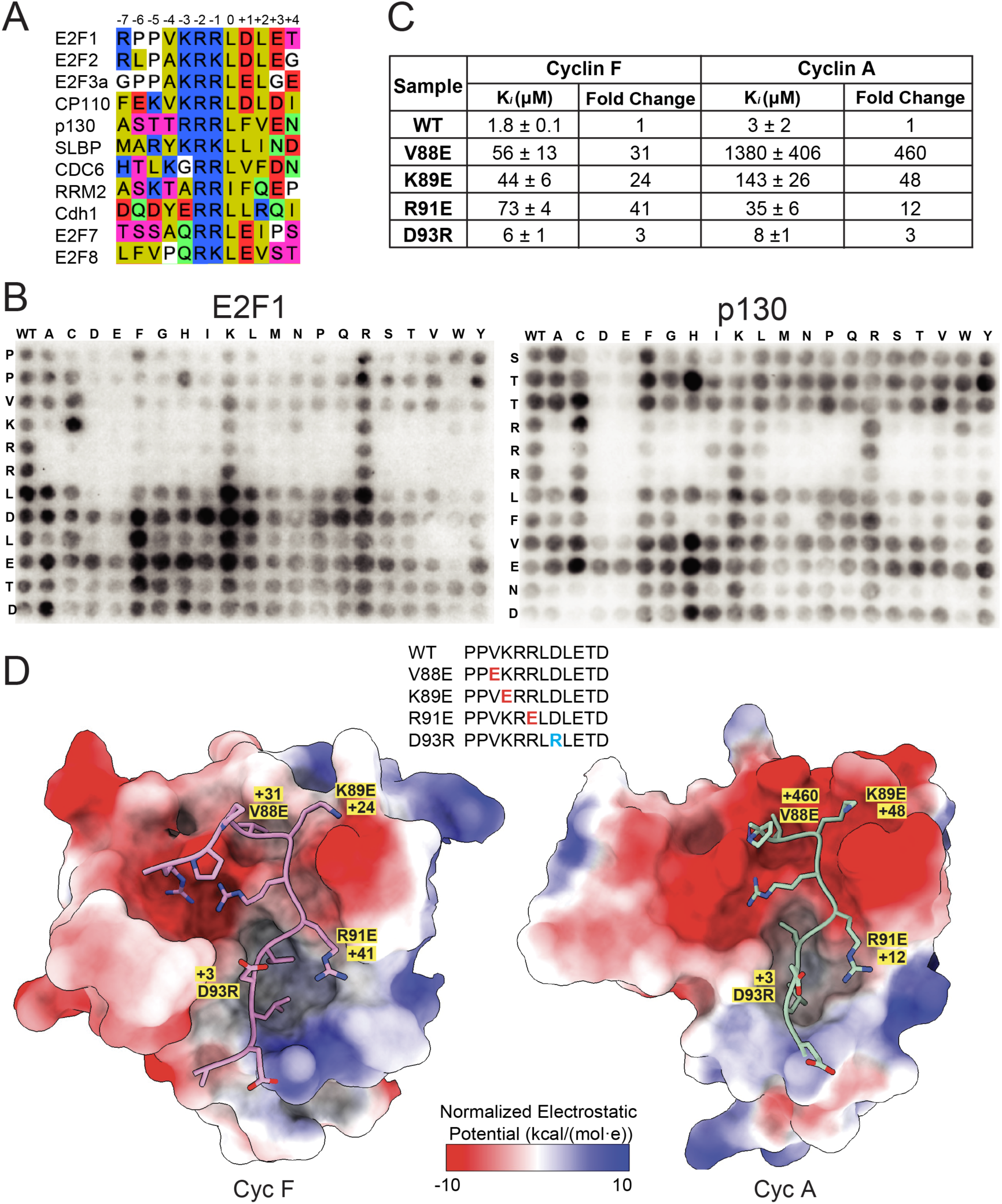
**A)** SPOT peptide array of E2F1 and p130 peptide probed for GST-Cyclin F (25-546)-GST-Skp1 binding with an anti-GST antibody. **B)** Sequence alignment of RxL motifs from Cyclin F substrates. **C)** Fluorescence polarization competition assay with E2F1 RxL mutant peptides. **D)** Coulombic electrostatic surface potential for Cyclin F and Cyclin A at the E2F1 peptide binding interface. Numbers represent the fold change increase KD associated with the mutation in E2F1 peptides (from panel C).

### Significance of Cyclin F specific interactions with E2F1 on association

These differences at the E2F1 interface suggest Cyclin F and Cyclin A have similar but non-identical substrate specificities. To measure the binding affinity of E2F1 peptide for Cyclin F and Cyclin A and test the importance of the non-conserved amino acids, we implemented a quantitative fluorescence polarization (FP) assay. As a probe, we used a TAMRA-labeled E2F1 peptide that contained the same sequence as the peptide used for EM studies. We found that both Cyclin A (Kd = 0.51 ± 0.05 μM) and Cyclin F-Skp1 (Kd = 0.89 ± 0.08 μM) bound the E2F1 peptide, with Cyclin A having slightly greater affinity (Figure 2C).

To quantitively assess the importance of the non-conserved amino acids that affect Cyclin F binding affinity for E2F1, we employed the same FP assay using Cyclin F mutants. We tested mutations to the extended loop, M323, which is C-terminal to the hydrophobic MYRIL sequence on α16, and R349 and Y350, which are located adjacent to the hydrophobic groove on α18 (Figure 2B). We also tested the mutation of the conserved hydrophobic MRYIL motif residues M310 and L313 for comparison. These indicated residues were mutated to alanine, mutant Cyclin F proteins were co-expressed with GST-Skp1 in SF9 insect cells, and the binding affinities for TAMRA-E2F1 peptide were determined (Figure 2D). Our data show that the hydrophobic MYRIL mutant (M309A/L313A) and the M323A mutant had the largest effect on E2F1 peptide binding; each mutation induced a a 9-fold decrease in affinity. The extended loop and R349/Y350 mutants had a 3-fold and 2-fold decrease in E2F1 peptide affinity, respectively.

To test the effect of the Cyclin F MRYIL and loop mutants on substrate binding in a cellular setting, we performed coimmunoprecipitation assays in HEK293T cells (Figures 2E and 2F). We could not detect sufficient signal from endogenous E2F1 protein, prompting us to instead overexpress E2F1 (Figure 2E). In addition, we probed for the endogenous Cyclin F substrates CP110 and CDC6 to assess their interactions under native expression levels (Figure 2F). As previously described, we found that the Cyclin F MRYIL mutant weakened the association with E2F1 (3), CP110, and CDC6 (6). The Cyclin F loop mutant showed a slightly weaker association for E2F1 compared to WT Cyclin F, and the weaker association was more apparent when probing for endogenously expressed proteins CP110 and CDC6 (Figure 2F). These data highlight residues outside of the conserved MRYIL motif that are important for Cyclin F to bind substrates.

### Significance of charged amino acids within the E2F1 RxL motif

We next aimed to test our hypothesis that substrate amino acids within and surrounding the RxL motif are important for Cyclin F and Cyclin A binding and contribute to substrate specificity. Comparing the RxL motif for various Cyclin F substrates reveals additional positively charged amino acids (K/R) in and near the RxL motif (Figure 3A), which suggests electrostatic interactions may be important drivers of affinity and specificity. To first map qualitatively the critical sequence requirements for Cyclin F substrate recruitment, we utilized SPOT peptide arrays in which E2F1 and p130 peptides were directly synthesized on a membrane (Figure 3B). Each spot contains a version of the E2F1 or p130 RxL sequence with a single substitution at each position in the peptide with each of the twenty standard amino acids. The array was probed with recombinant GST-Cyclin F (25-546)-GST-Skp1 heterodimer and an anti-GST antibody. Although there was some variability in the signal that precludes rigorous quantitative analysis, there were some clear trends, especially with respect to the effects of charged amino acids. Defining the 0 position as the L in the RxL motif, we observed that positive amino acids (K/R) in the -3, -2, and -1 positions were necessary for a robust interaction and that introducing a negatively charged amino acid (D/E) in the -6, -5, -4, -3, -2, -1, and 0 in E2F1 or the -6, -5, -4, -3, -2, -1, 0, and +1 position of p130 inhibited binding. These results further suggest that electrostatics play an important role in Cyclin F recognizing its substrates, where a positive charge in the -3, -2 and -1 enabled Cyclin F binding while a negative charge in the -6, -5, -4, -3, -2, -1, and 0 for E2F1 decreases affinity for Cyclin F.

To validate and test more quantitatively the significance of electrostatics for the recruitment of substrates by Cyclin F and Cyclin A, we implemented competitive binding FP experiments using peptides in which the charged residues surrounding the RxL motif in E2F1 were substituted for the opposite charge. The results correlated well with the E2F1 SPOT array. Specifically, a negatively charged glutamic acid mutation in the -1 (R91E), -3 (K89E), and -4 (V88E) positions decreased binding affinity for Cyclin F by 41-fold, 24-fold, and 31-fold, respectively (Figure 3C and 3D). A positive charge in the +1 (D93R) position had a more subtle 3-fold decrease in affinity for Cyclin F. The same competitive binding FP experiment was performed with Cyclin A and revealed some notable differences. Introducing a negative charge in the -1 (R91E), -3 (K89E), and -4 positions (V88E) resulted in a larger decrease in affinity by 12-fold, 48-fold, and 460-fold, respectively, while a positive charge in the +1 (D93R) position had a 3-fold decrease in affinity for Cyclin A. These results suggest that a negative charge in the -3 and -4 positions of the substrate is less tolerated by Cyclin A, a negative charge in the -1 position is less tolerated by Cyclin F, and a positive charge in the +1 position has a minor effect on both Cyclin F and Cyclin A binding.

## Discussion

Our structural data reveals the architecture of the Skp1-Cyclin F complex and how it recognizes the RxL critical for substrate binding. We find that the CDK blocking domain (CBD) is a novel structural feature that further distinguishes Cyclin F from canonical cyclins. The precise functional role of the CBD remains to be fully elucidated. One study suggests that the CBD-cyclin box domain interface may create a secondary substrate binding site on Cyclin F, potentially coordinating a phosphorylated Thr/Ser residue on Exo1 (8). Interestingly, many other known Cyclin F substrates, including E2F1 and RRM2 (5, 6), contain a putative CDK phosphorylation site (T/SP) upstream of the RxL motif, positioning them for interaction with this secondary binding site. Support for this hypothesis comes from AlphaFold models (Supplemental Figure 10), which demonstrate that residues Y287 and T283 of the cyclin box, alongside residues 162-164 of the CBD, could form a phosphate-binding pocket. While further experimental validation is required to confirm this interaction, we propose that in general, the CBD has great potential for facilitating Cyclin F interactions with other protein regulators and substrates.

The CBD may also play a critical role in substrate positioning (Figure 4). As a substrate recognition protein for the SCF E3 ubiquitin ligase, Cyclin F binds substrates and positions them for ubiquitylation. The network of hydrophobic interactions that connects the F box domain through the CBD to the cyclin domain suggests that the orientation of the MRYIL site relative to the rest of the ligase is fixed. The CBD may therefore facilitate ubiquitylation by constraining the steric freedom of bound substrates or play a role in dictating which lysines in bound substrates are ubiquitylated by holding them in a specific orientation relative to the SCF. We also suspect that the CBD plays an important scaffolding role by stabilizing the cyclin box domain. We were unsuccessful in attempts to express the cyclin box domain of Cyclin F alone, suggesting that CBD is required for the proper formation of the cyclin box structure.

**Figure 4.**
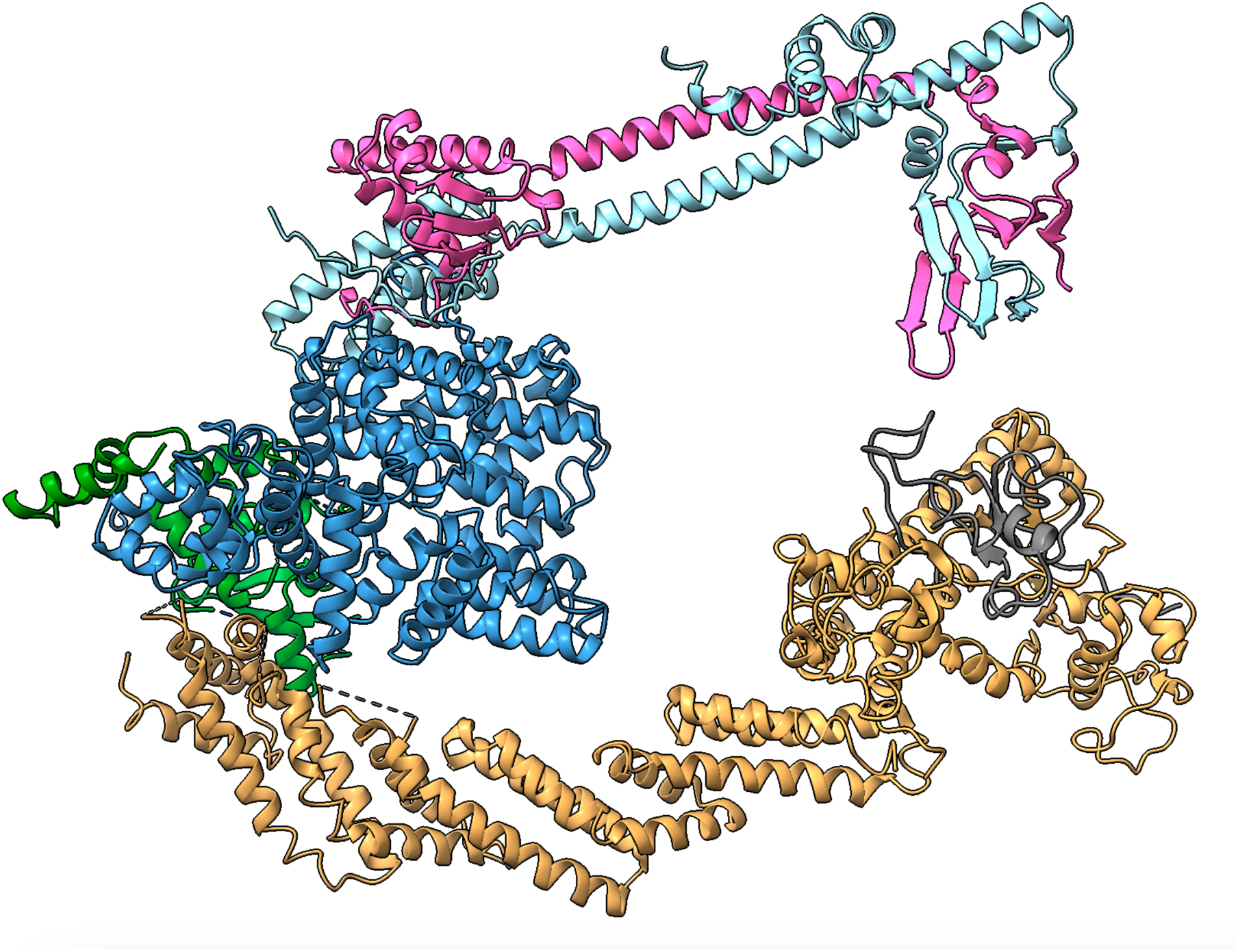
Model of SCF^CyclinF^ E3 ubiquitin ligase complex bound to E2F1-DP1. E2F1 (pink), DP1 (light blue) is positioned by Cyclin F (blue). Skp1 (green), Cul1 (yellow), and Rbx1 (grey) are shown.

We have identified important differences in how Cyclin F and Cyclin A bind substrates that contain RxL motifs. We found that changing the charge on the E2F1 peptide affects Cyclin F and Cyclin A differently; V88E and K89E decrease the affinity for Cyclin A more, while R91E decreases the affinity for Cyclin F more (Figure 3C). Structural data suggest that these differences are due to non-conserved amino acids in the cyclin domain near the RxL-binding interface that result in a difference in the electrostatic environment. For example, Cyclin F has an extended loop (D504-T511) adjacent to the MRYIL site that affects substrate binding when mutated (Figure 2D-2F), and Cyclin A has a larger electronegative hotspot at the C-terminal end of helix 1 (Figure 3D). Targeting cyclin proteins for therapeutics has been discussed in the literature as an alternative to targeting CDKs (23–25). The difference in structure and substrate interactions can aid in the development of Cyclin F or Cyclin A-specific therapeutics.

## Methods

### Protein expression and purification

Strep-Cyclin F (1-546 and 1-636) and GST-Cyclin F (25-546) fusion proteins were co-expressed with GST-Skp1 (1-163) in SF9 cells via baculovirus infection. Cell pellets were harvested after 72 h of growth in suspension at 27 °C and lysed by sonication in 25 mM Tris pH 8.0, 500 mM NaCl, 5 mM DTT, and 5% glycerol. Cyclin F-Skp1 complexes were purified using GST-affinity purification. After removal of the affinity tags through TEV protease cleavage, purified complexes were concentrated and further purified through size-exclusion chromatography using a Cytiva Superdex 200 10/300 increase column. The final buffer contained 25 mM Tris pH 8.0, 400 mM NaCl, and 3 mM DTT.

### Single-Particle Cryo-EM analysis

Purified Cyclin F (1-636)-Skp1 complex was incubated with a synthetic E2F1 peptide (84-GRPPVKRRLDLETDHQ-99) at a 2:1 molar ratio and the E2F1-Cyclin F-Skp1 complex was isolated using a Cytiva Superdex 200 10/300 increase column in 150 mM NaCl, 25 mM Tris HCl, and 1 mM DTT at pH 8.0. The protein complex was set on previously glow-discharged Au-ultraFoil R1.2/1.3 400 mesh grids at a concentration of 1 mg/mL. 3.5 μL of sample were fast-plunged into liquid ethane using a ThermoFisher Scientific (TFS) Vitrobot Mark IV set at 100% humidity and 4 °C and blotted for 1.5 s.

Initial grids screening was performed at UCSC Biomolecular cryoEM Facility using a TFS Glacios 200 kV coupled to a Gatan K2 Summit. The top-selected grids were collected at the UCSF CryoEM Facility using a TFS Krios G2 using Gatan K3 coupled to a Biocontinuum energy filter (10 eV). The 7,611 images were acquired at an accelerating voltage of 300 kV and nominal magnification of x105,000, giving a calibrated pixel size of 0.835 Å, exposure time of 0.57 s and total fluence of 45.8 e/Å^2^.

Acquired data was processed using cryoSPARC v4.4.1 workflow (Supplemental Figure 2), including motion correction and contrast transfer function (CTF) estimation. Particles were picked using a bias-free blob picker with an 80 Å minimum and 120 Å maximum particle diameter. Accepted 7,781,999 particles were extracted with a box size of 256 pixels. Extracted particles were subject to five rounds of 2D classification and selected particles were used for Ab initio reconstruction followed by non-uniform refinement which resulted in a map that encompassed the full complex. To improve resolution, 3D classification, followed by non-uniform refinement was conducted to generate a final map with an overall resolution of 3.47 Å and local resolution spanning 3.0-4.0 Å.

### Model building and structural analysis

The initial model for E2F1 peptide-Cyclin F-Skp1 complex was generated by AlphaFold and docked into the map using Coot v0.9.4 (35). Further modeling, coordinate refinement, and energy minimization were performed using Coot v0.9.4 and Phenix v. 1.20.1-4487 (36, 37). Validation was performed using FSC-Q (38) and model-to-map criteria (39) (Table 1). The coulombic electrostatic potential (ESP) calculation was performed using ChimeraX v1.8.

**Table 1:** CryoEM data collection, single-particle reconstruction maps, and model statistics.

|  |  |
| --- | --- |
| **Data collection information** | **UCSF – Krios G2** |
| Nominal magnification | 105,000x |
| Voltage (kV) | 300 |
| Electron dose (e<sup>-</sup>/Å<sup>2</sup>) | 45.8 |
| Physical pixel size (Å) | 0.835 |
| Movies amount | 7,611 |
| Defocus average and range (μm) | 1.5 (0.5-2.5) |
| Frames | 79 |
| **Single-particle reconstruction information** | **EMD-45413** |
| Initial particles picked | 7,781,999 |
| Particles in the final map | 103,503 |
| Initial model used | Ab initio |
| Symmetry imposed | C1 |
| Map overall resolution - FSC<sub>0.143</sub> (Å) | 3.47 |
| Map resolution range (Å) | 3.0-4.0 |
| **Built model information** | **PDB 9CB3** |
| Chains | 3 |
| Atoms (non-H) | 4,995 (H: 0) |
| Protein residues | 672 |
| Ligands | 0 |
| **Bonds (RMSD)** |  |
| Length (Å, # > 4σ) | 0.015 (3) |
| Angles (°, # > 4σ) | 0.65 (19) |
| **Mean B-factor (Å<sup>2</sup>)** | 4995/0 |
| Protein | 30.00/183.57/105.62 |
| (min/max/mean) |  |
| **MolProbity score** | 2.08 |
| Clashscore | 12.53 |
| Rotamers outliers (%) | 1.46 |
| **Ramachandran plot** |  |
| Outliers | 0.01 |
| Allowed (%) | 5.01 |
| Favored (%) | 94.99 |
| **Rama-Z (Ramachandran plot Z-score, RMSD)** |  |
| Whole (N = 661) | 0.03 (0.34) |
| Helix (N = 451) | 0.65 (0.26) |
| Sheet (N = 0) | - (-) |
| Loop (N = 210) | 1.27 (0.44) |
| **Rotamer outliers (%)** | 1.46 |
| Cβ outliers (%) | 0.15 |
| **Peptide plane (%)** |  |
| Cis proline/general | 4.5/0.0 |
| Twisted proline/general | 0.0/0.0 |
| **CaBLAM outliers (%)** | 2.78 |
| **Occupancy** |  |
| Mean | 1.00 |
| occ = 1 (%) | 99.56 |
| 0 < occ < 1 (%) | 0.44 |
| occ > 1 (%) | 0.00 |
| **Model vs. Data** |  |
| Mean CC for ligands | -- |
| CC (peaks) | 0.49 |
| CC (volume) | 0.65 |
| CC (box) | 0.57 |
| CC (mask) | 0.68 |

The maps and model were deposited in the Electron Microscopy Databank and Protein Data Bank, respectively, with accession numbers EMD-45413 and PDB ID 9CB3. Raw data and processed files are publicly available at EMPIAR-12145. Data collection and model refinement parameters are reported in Table 1. Figures were generated using UCSF Chimera-X (40).

### Fluorescence polarization (FP) binding experiments

TAMRA-labeled E2F1^84-99^ peptide was mixed at 10 nM with strep-Cyclin F (1-546)-GST-Skp1 (1-163) (S1F) in a buffer containing 25 mM Tris pH 8.0, 150 mM NaCl, 1 mM DTT, and 0.1% (v/v) Tween-20. Samples were incubated for 1 hour at 4 °C and 40 μL of the reaction was used for the measurement in a 384-well plate. For the competition experiments, wild-type or mutant, unlabeled E2F1 peptide were mixed in a buffer containing 10 nM TAMRA-E2F1^84-99^ and 800 nM S1F or 500 nM GST-Cyclin A (173-432) in 25 mM Tris pH 8.0, 150 mM NaCl, 1 mM DTT, and 0.1% (v/v) Tween-20. Samples were incubated for 1 hour at 4 °C and 40 μL of the reaction was used for the measurement in a 384-well plate. FP measurements were made in triplicate, using a Perkin-Elmer EnVision plate reader. The KD, IC50, and Ki values were determined using Prism 8 (Version 8.4.3).

### Spot blot peptide array

RxL containing E2F1 and p130 12-mer peptide arrays were purchased from Kinexus. Synthetic peptides on TOTD membranes were hydrated in 100% methanol while shaking for 10 minutes, followed by 3 washes in 20 mM Tris pH 7.6, 150 mM NaCl (Tris-buffered saline, TBS) for 5 minutes each. Membranes were transferred into blocking buffer (TBS-T with 5% FBS) for 2 hours at room temperature followed by one wash in TBS-T for 5 minutes. GST-Cyclin F (25-546)-GST-Skp1 (1-163) protein complex was diluted to 1 μg/mL in TBS-T, and membranes were incubated overnight at 4 °C. Membranes were washed 3 times in TBS-T for 5 minutes, and membranes were incubated in HRP-conjugated anti-GST antibody in blocking buffer for 2 hours at room temperature. Membranes were washed three times with TBS-T for 5 minutes followed by one wash in TBS for 5 minutes at room temperature. Excess buffer was removed and chemiluminescence was measured using SuperSignal West Pico Plus substrate.

### Immunoprecipitation

HEK293T cells were transfected with tagged protein constructs for 48 hours prior to harvesting for lysis and subsequent immunoprecipitation. Briefly, transfected cells were washed three times with ice-cold 1x PBS and harvested by scraping. The cells were then pelleted by centrifugation at 1,000xg for 5 minutes at 4 °C. Washed cell pellets were lysed using NETN lysis buffer (20 mM Tris pH 8.0, 100 mM NaCl, 0.5 mM EDTA, 0.5% NP40) with the addition of 10 μg/ml of Leupeptin, 10 μg/ml of aprotonin, 10 μg/ml of pepstatin A, 1 mM of AEBSF, 1 mM of sodium fluoride, 1 mM of sodium orthovanadate for 15 minutes on ice. Cell lysates were spun down to remove debris for 15 min at maximum speed in a refrigerated microcentrifuge at 4 °C. Protein concentration was determined by the Bradford Assay. Prior to immunoprecipitation, 50 μl of anti-Flag M2 affinity gel (Sigma-Aldrich, Cat. #F2426) was washed 2x of cold PBST and 1x of NETN lysis buffer at 4 °C while rotating for 5 minutes. 10% of total protein was reserved as input. The remaining lysate was added to the washed anti-Flag M2 affinity gel and immunoprecipitated for 1 hour at 4 °C while rotating. Then anti-Flag M2 affinity gel was washed 3x with NETN lysis buffer and resuspended in 50 μl of 2x Laemmli sample buffer. Samples were heated to 95°C for 5 minutes prior to analysis by immunoblotting.

### Data Accessibility

The cryoEM density map has been deposited in the Electron Microscopy Data Bank under accession code EMD-45413 and coordinates have been deposited in the Protein Data Bank under the accession code PDB ID 9CB3. Raw data were deposited in the EMPIAR (EMPIAR-12145).

## Acknowledgments

This work was supported by NIH grants to S. Rubin (P01CA254867 and R35GM145255) and to M. Emanuele (R35GM153250). We thank Dr. David Bulkley and the UCSF CryoEM Facility for data collection. The authors also acknowledge the Biomolecular cryo-Electron Microscopy Facility at the Department of Chemistry and Biochemistry of the University of California - Santa Cruz (RRID: SCR_021755) for scientific and technical assistance (NIH High-End Instrumentation program, S10OD02509). Molecular graphics and analyses performed with UCSF ChimeraX, developed by the Resource for Biocomputing, Visualization, and Informatics at the University of California, San Francisco, with support from National Institutes of Health R01-GM129325 and the Office of Cyber Infrastructure and Computational Biology, National Institute of Allergy and Infectious Diseases.

---

**Supplemental Figure 1.**
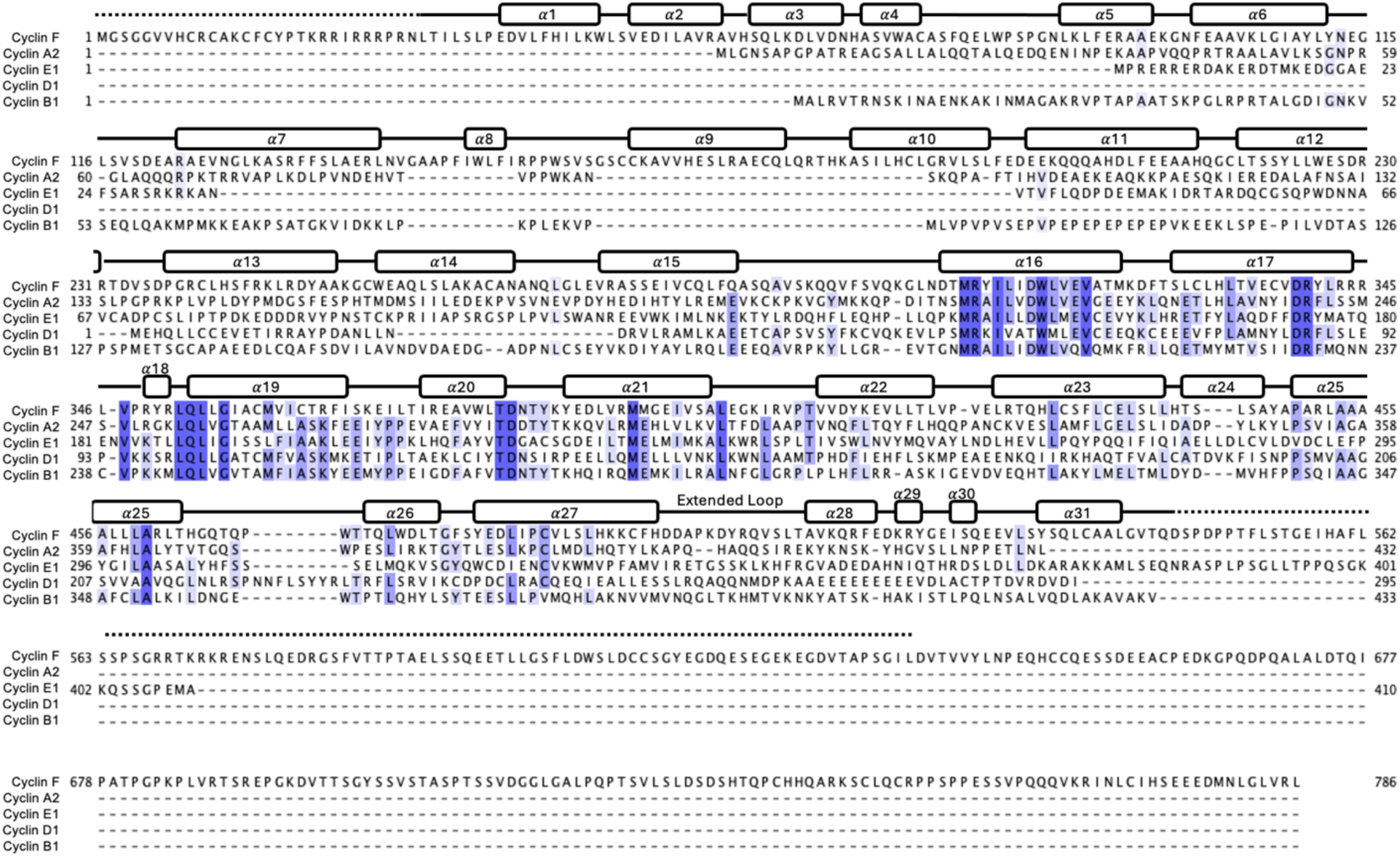
Sequence alignment of canonical cell-cycle cyclins and Cyclin F. Secondary structure of Cyclin F is shown. The dotted line appears over sequences not modeled from cryo-EM data but present in the protein construct. Sequences were aligned with ClustalWS with default settings and shading represents amino acids that are above the conservation threshold of 60%.

**Supplemental Figure 2.**
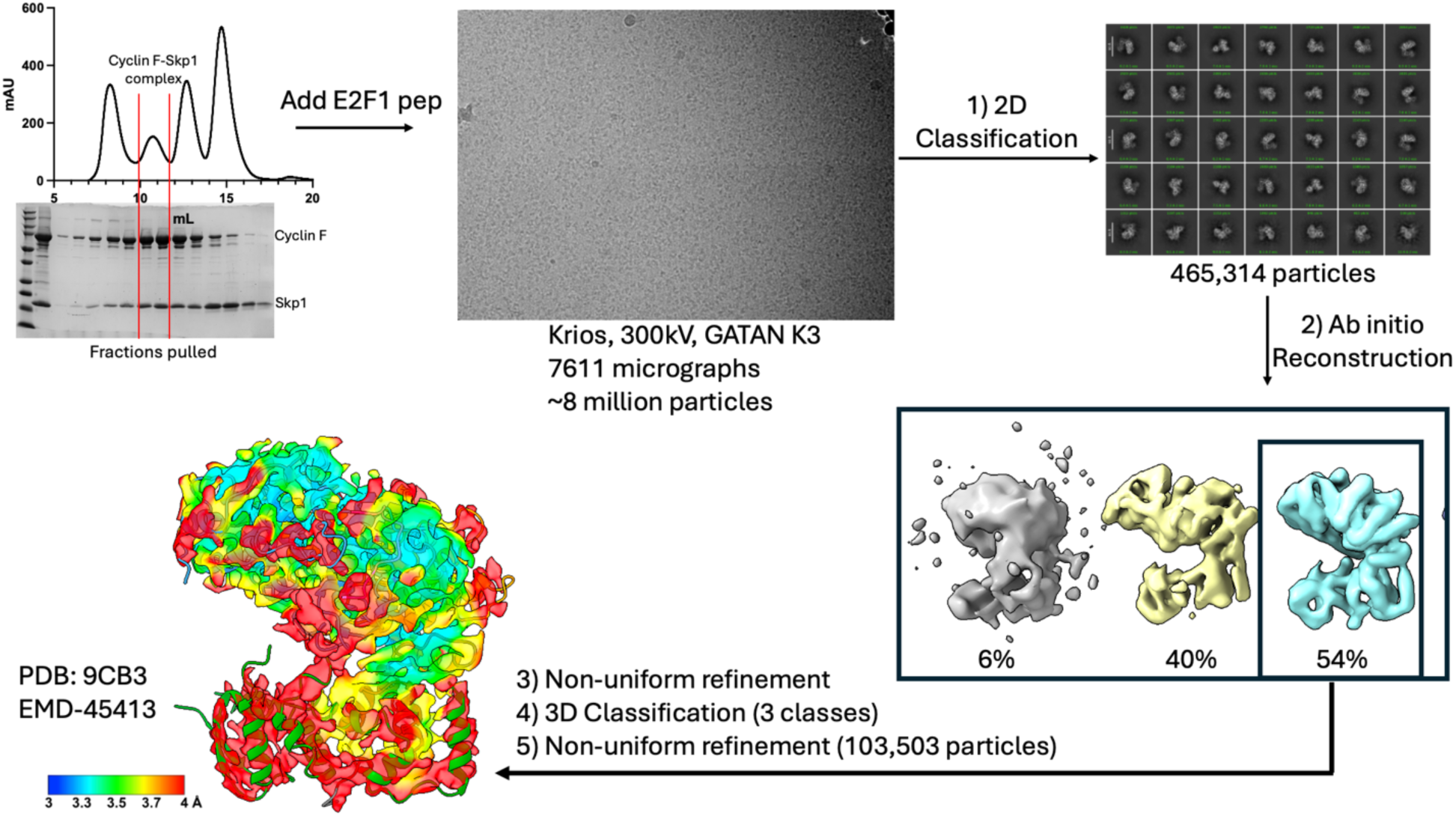
Single-particle cryo-EM data processing workflow.

**Supplemental Figure 3.**
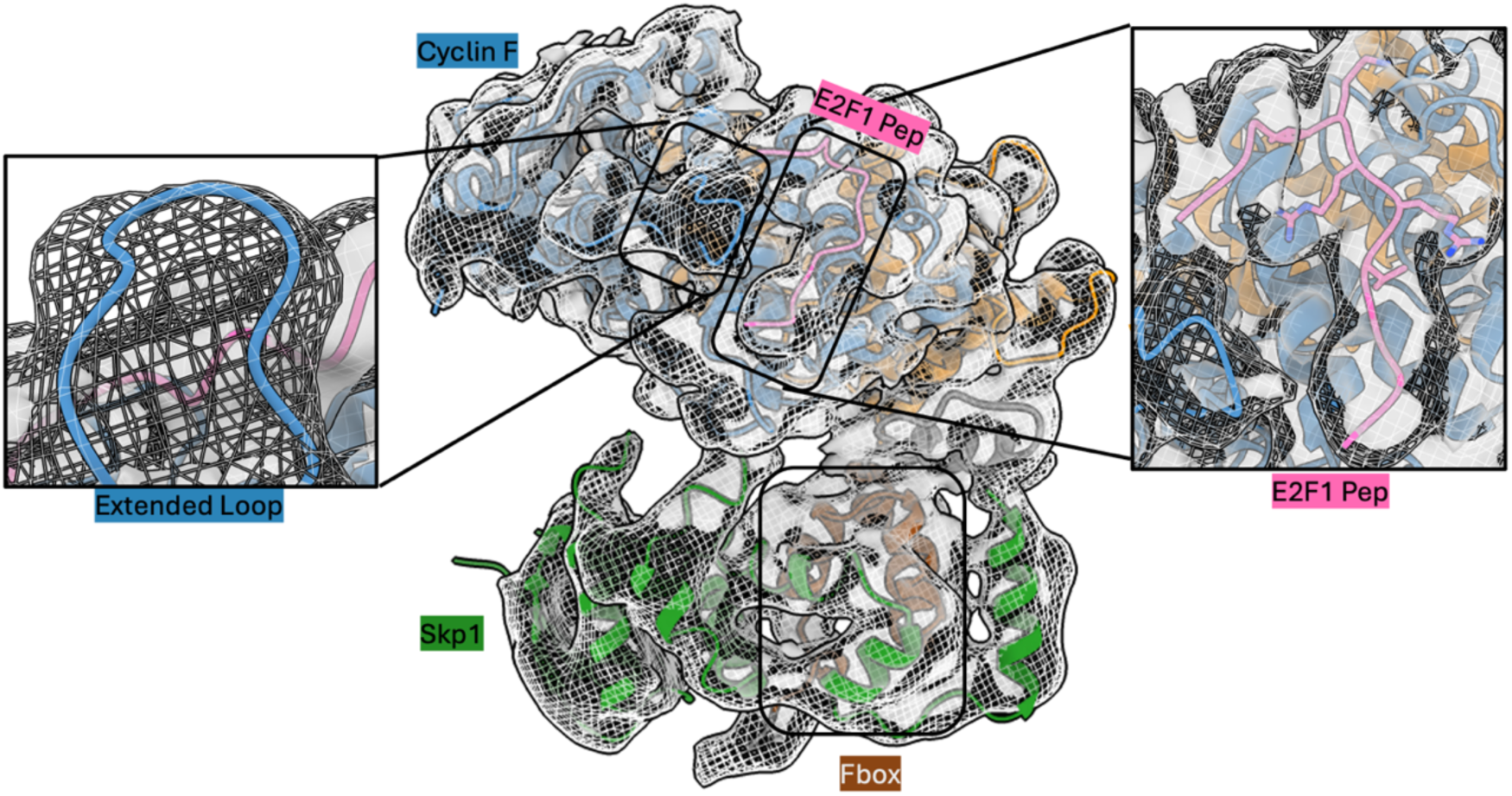
Model of E2F1 peptide-Cyclin F-Skp1 builds within electron density. The mesh surface indicates an unsharpened electron density map, and the grey surface indicates a sharpened electron density map.

**Supplemental Figure 4.**
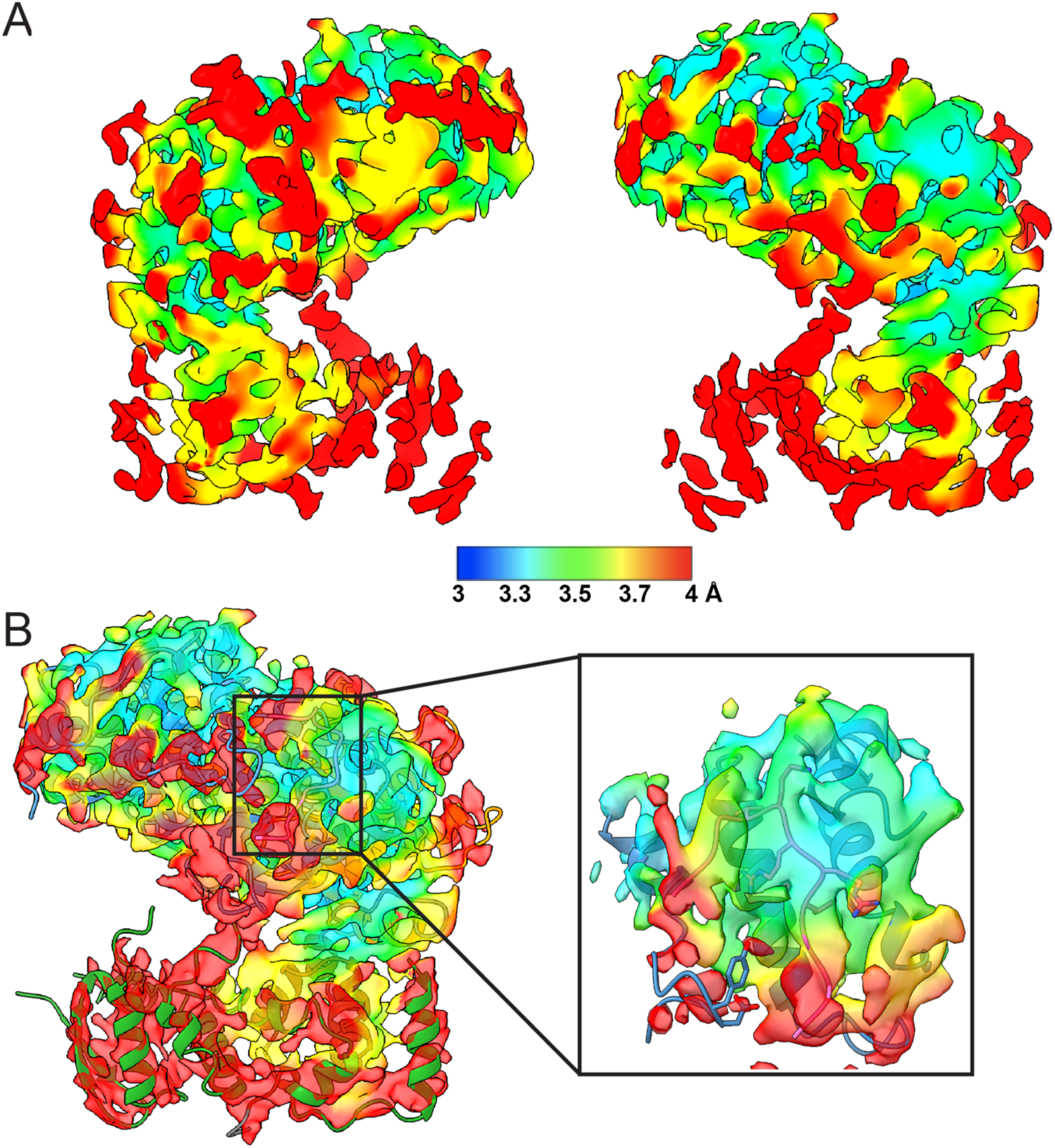
**A)** Local resolution of electron density map. **B)** Local resolution at the E2F1 binding interface. The model for the E2F1 peptide was built at a contour level of 0.212 and the extended loop was built at a contour level of 0.153.

**Supplemental Figure 5.**
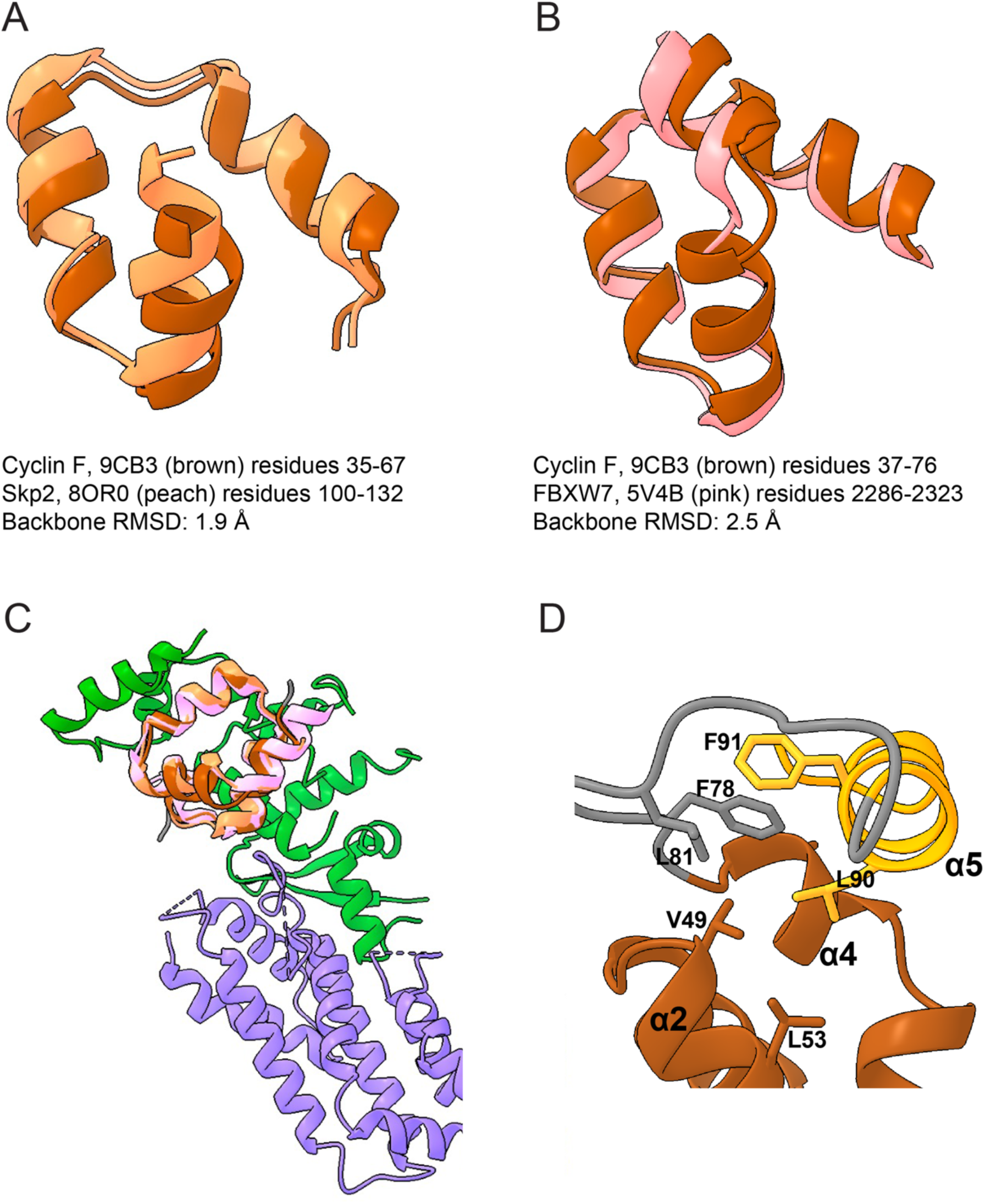
F box backbone RMSD calculations between **A)** Cyclin F and Skp2 and **B)** Cyclin F and FBXW7. **C)** Structural alignment of F box domains (Cyclin F, Skp2, and FBXW7), Skp1 (green, 9cb3), and Cul1 (purple, 8OR0). **D)** Interactions bridging the F box to the CBD in Cyclin F.

**Supplemental Figure 6.**
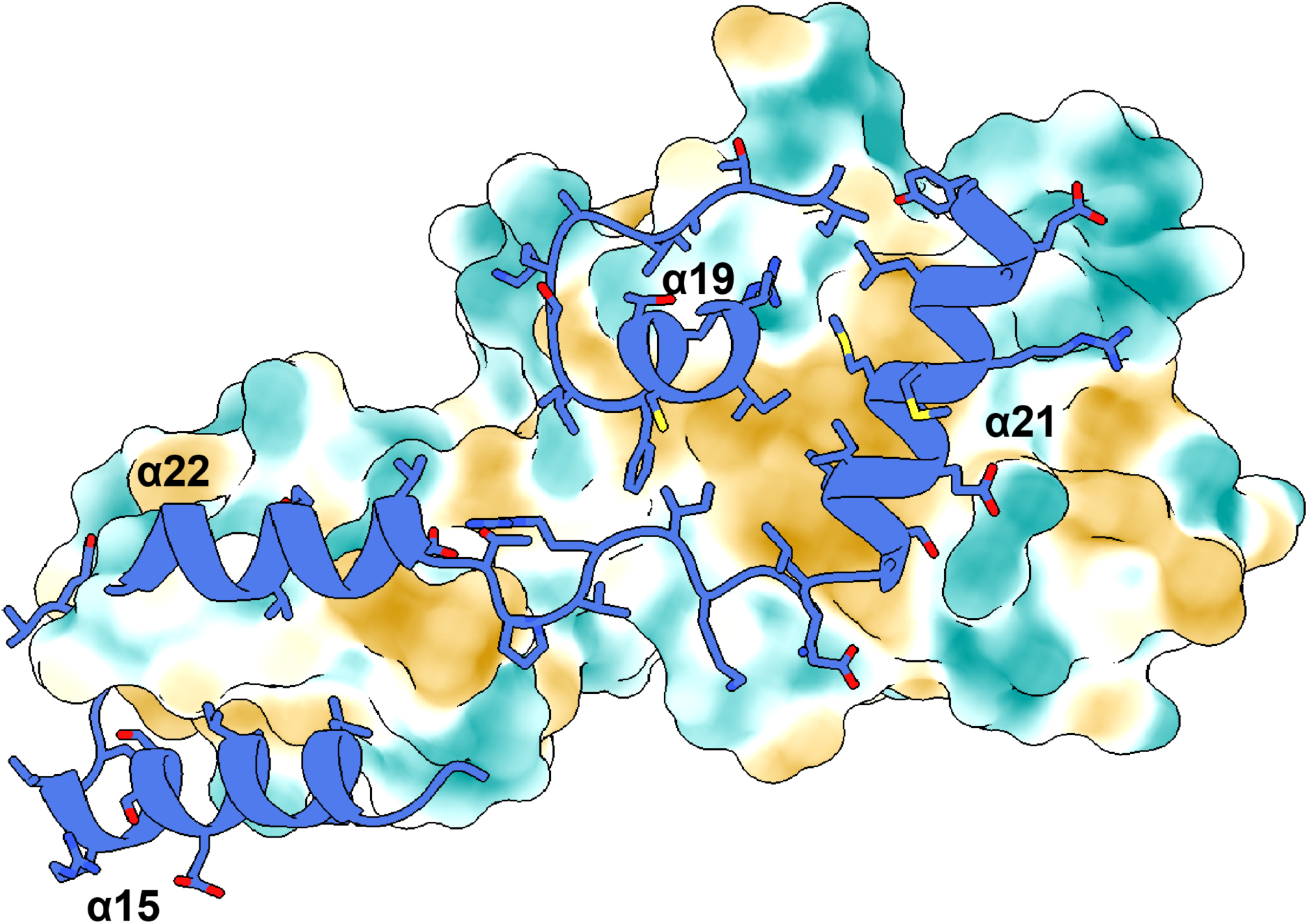
CBD with hydrophobic surface area (brown is hydrophobic surface and turquoise is hydrophilic surface). The cyclin box at the interface is shown as ribbons, and the helices that contact the CBD are indicated.

**Supplemental Figure 7.**
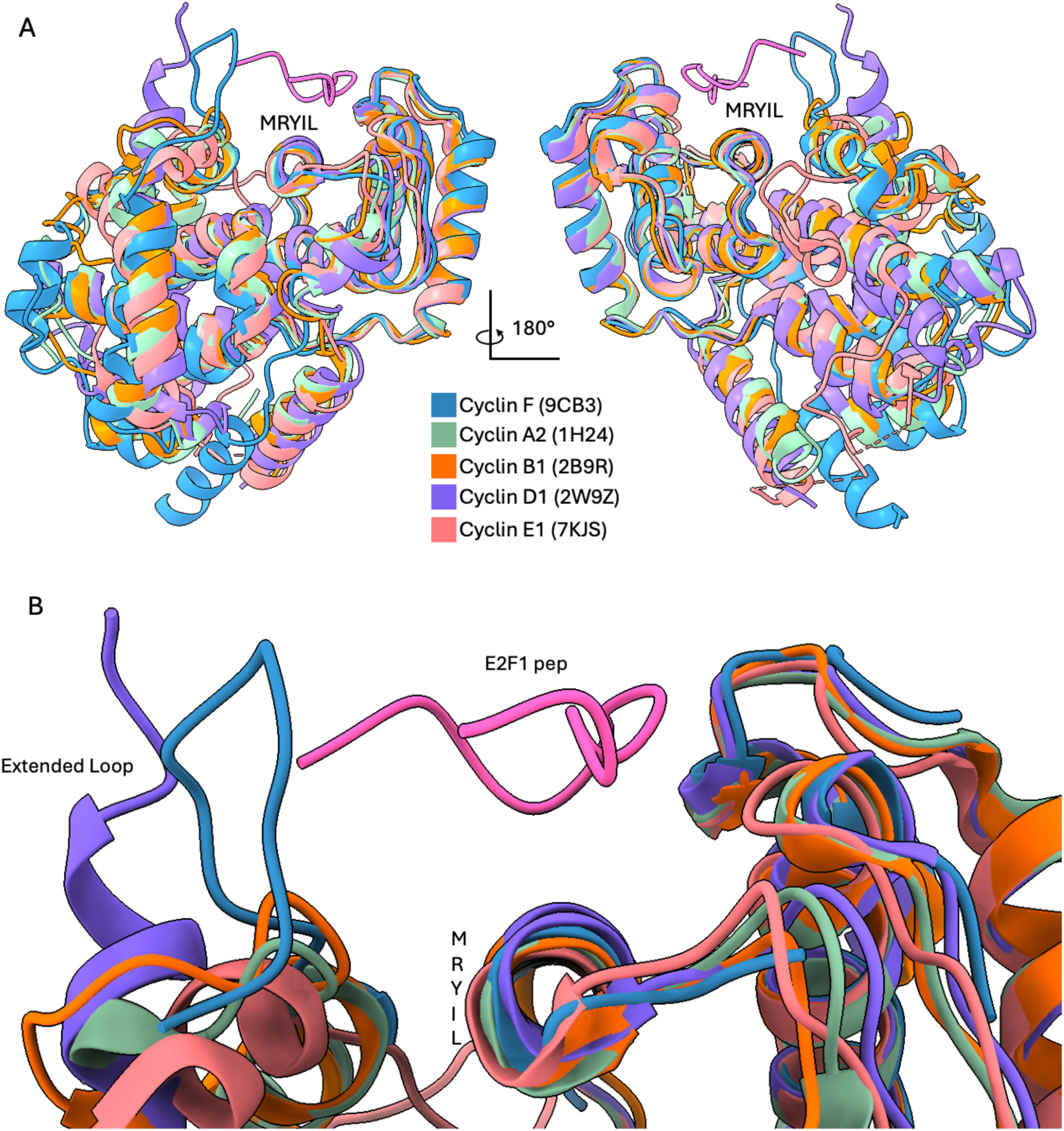
**A)** Structural alignment of the Cyclin F cyclin box domain with canonical cell-cycle cyclin proteins. The E2F1 RxL motif peptide is shown in pink above the MRYIL site. **B)** Close-up view of the cyclin box binding interface with E2F1.

**Supplemental Figure 8.**
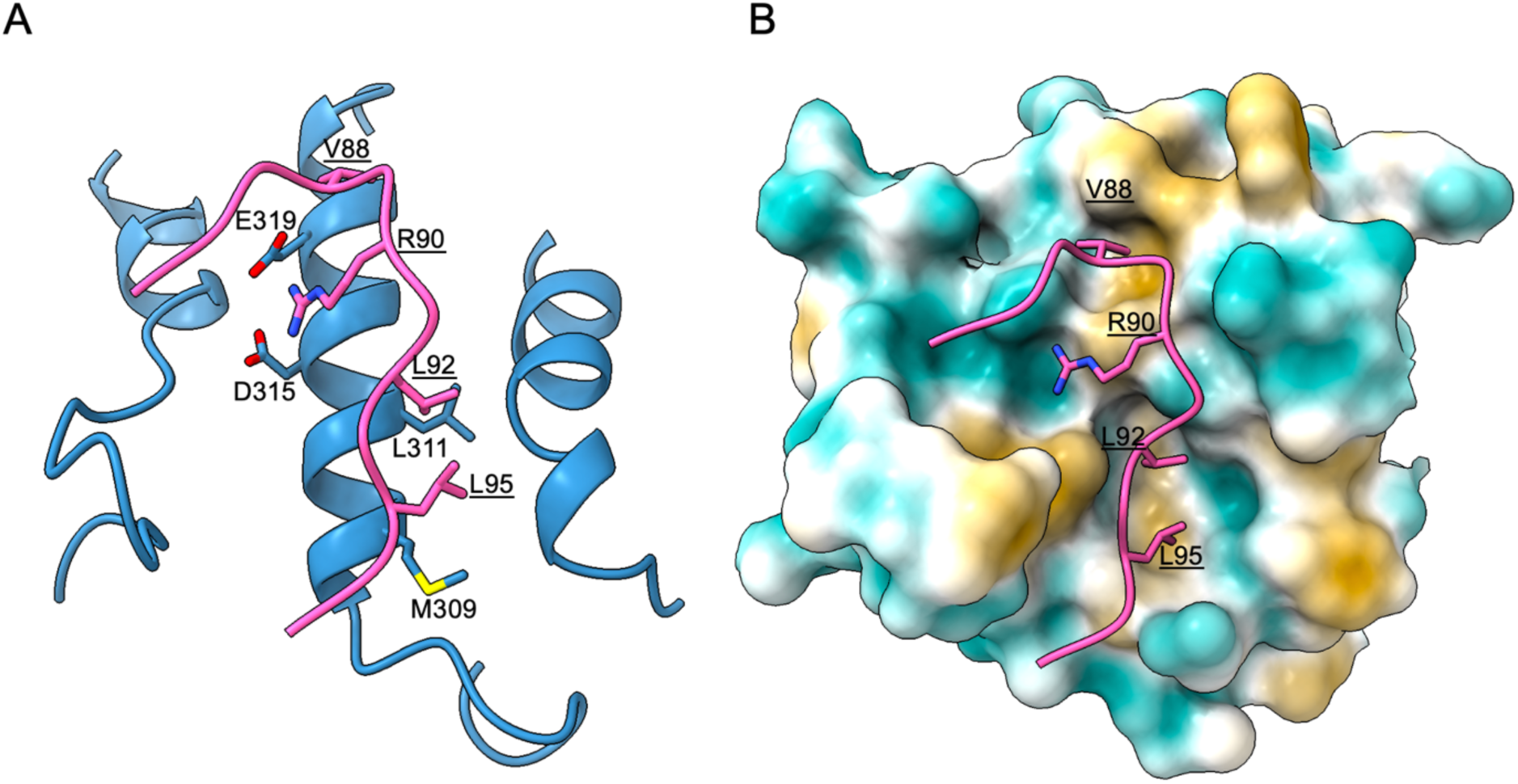
Cyclin F-E2F1 peptide binding interface. The surface area of Cyclin F is shown with brown indicating hydrophobic areas and turquoise indicating hydrophilic areas.

**Supplemental Figure 9.**
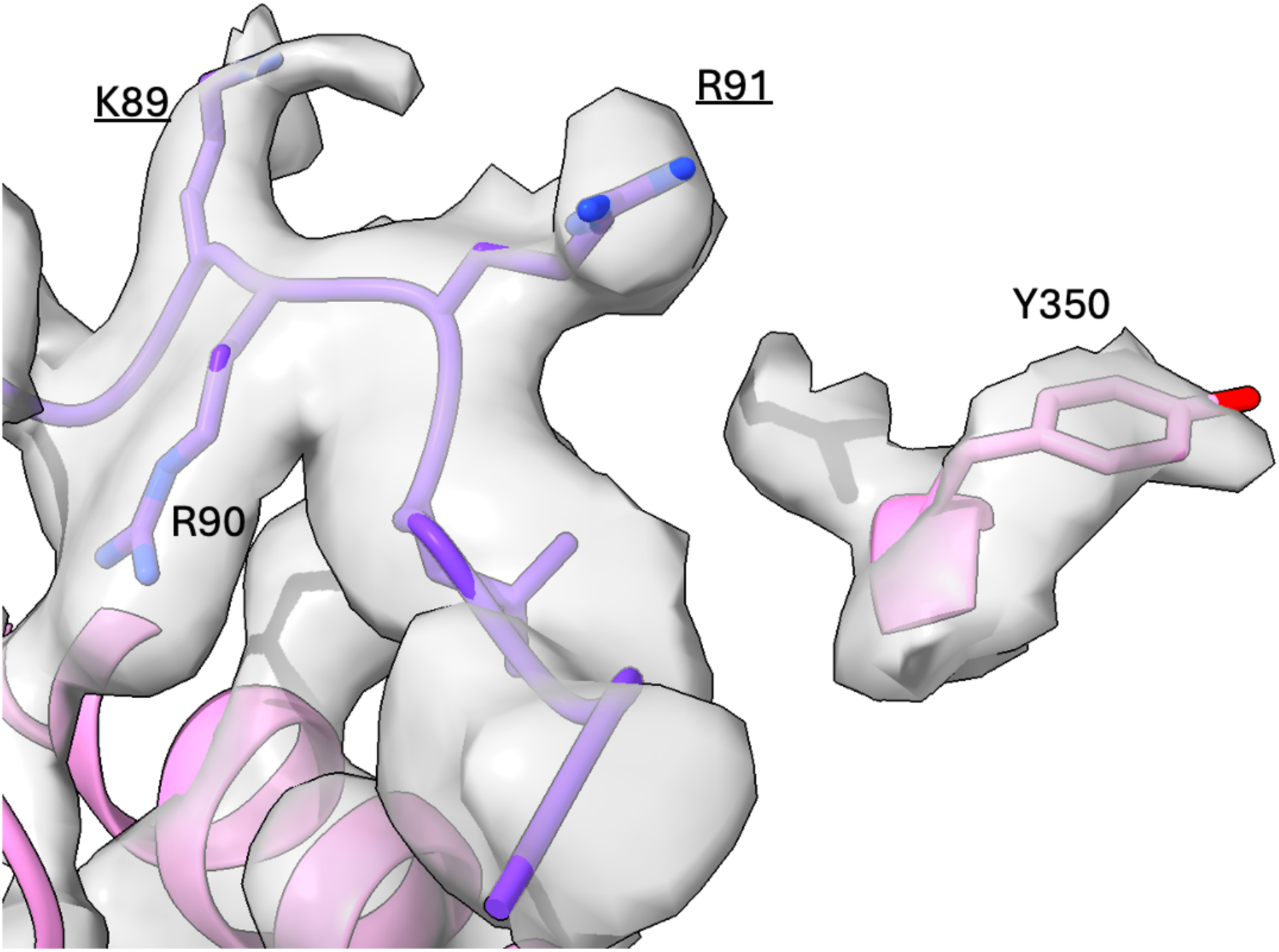
Electron density around Y350 at contour level 0.212.

**Supplemental Figure 10.**
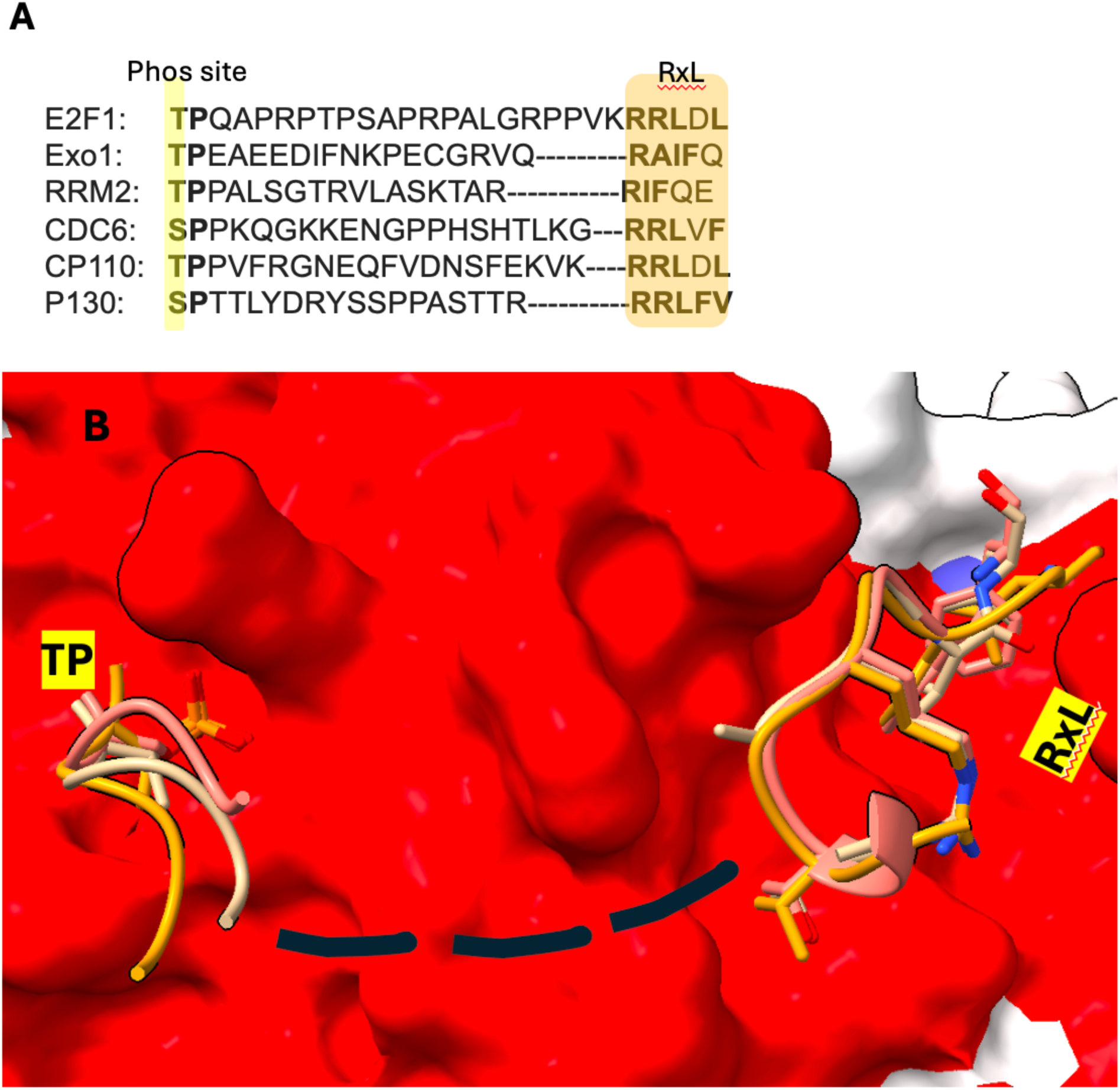
**A)** Putative phosphorylation site upstream of RxL motif for Cyclin F substrates. **B)** AlphaFold3 model of phosphorylated Cyclin F substrates (E2F1 (orange), Exo1 (salmon), and RRM2 (beige)) bound to Cyclin F (red surface). The model was made on the AlphaFold Server (https://golgi.sandbox.google.com/) with Cyclin F (1-547) and phosphorylated peptide substrates for E2F1 (FApTPQAPRPTPSAPRPALGRPPVKRRLDLE), p130 (TpSPTTLYDRYSSPPASTTRRRLFVE), RRM2 (pTPPALSGTRVLASKTARRIFQ), and Exo1 (NIQLpTPEAEEDIFNKPECGRVQRAIFQ) as input sequences.

